# *Enterococcus faecalis* OG1RF Evolution at Low pH Selects Fusidate-sensitive Mutants in Elongation Factor G and at High pH Selects Defects in Phosphate Transport

**DOI:** 10.1101/2023.03.22.533894

**Authors:** Bailey A. Fitzgerald, Ayman Wadud, Zachary Slimak, Joan L. Slonczewski

## Abstract

*Enterococcus* bacteria inhabit human and soil environments that show a wide range of pH. Strains include commensals as well as antibiotic-resistant pathogens. We investigated adaptation to pH stress in *E. faecalis* OG1RF by conducting experimental evolution in acid (pH 4.8), neutral pH (pH 7.0), and base (pH 9.0). Serial planktonic culture was performed for 500 generations, and in high-pH biofilm culture for four serial bead transfers. Nearly all mutations led to nonsynonomous codons, indicating adaptive selection. All acid-adapted clones from planktonic culture showed a mutation in *fusA* (encoding elongation factor G). The acid-adapted *fusA* mutants had a tradeoff of decreased resistance to fusidic acid (fusidate). All base-adapted clones from planktonic cultures, and some from biofilm-adapted cultures, showed mutations affecting the Pst phosphate ABC transporter (*pstA, pstB, pstB2, pstC*) and *pyrR* (pyrimidine biosynthesis regulator/uracil phosphoribosyltransferase). Biofilm culture produced small-size colonies on brain-heart infusion agar; these variants each contained a single mutation in *pstB2*, *pstC*, or *pyrR*. The *pst* and *pyrR* mutants outgrew the ancestral strain at pH 9.2, with a tradeoff of lower growth at pH 4.8. Additional genes that had a mutation in multiple clones evolved at high pH (but not at low pH) include *oppBCDF* (oligopeptide ABC transporter), *ccpA* (catabolite control protein A), and *ftsZ* (septation protein). Overall, experimental evolution of *E. faecalis* showed strong pH dependence, favoring fusidate-sensitive elongation factor G modification at low pH and loss of phosphate transport genes at high pH.

**IMPORTANCE:** *E. faecalis* bacteria are found in dental biofilms where they experience low pH as a result of fermentative metabolism. Thus the effect of pH on antibiotic resistance has clinical importance. In endodontal infections, enterococci can resist calcium hydroxide therapy that generates extreme high pH. In other environments such as soil and plant rhizosphere, enterococci experience acidification associated with climate change. Thus the pH modulation of natural selection in enterococci is important for human health as well as for understanding soil environments.

## INTRODUCTION

*Enterococcus faecalis* are gram-positive fermenter bacteria found in a wide range of habitats, including dental biofilms and the gastrointestinal tract of humans (1–4) as well as soil and plants (5). Pathogenic strains cause a variety of infections throughout the body, which are of special concern in hospitals due to the acquired vancomycin-resistance in certain strains and an intrinsic antibiotic-resistance to numerous other antibiotics (6). *Enterococcus* species are known for their ability to tolerate a wide range of conditions such as salinity, temperature, and pH (6–9). Survival at low pH is of importance for enteric infections, and at high pH is relevant to dental biofilms that are treated with calcium hydroxide pastes (10, 11). Multidrug-resistant enterococci are isolated from environmental sources such as soil (12) and the digestive microbiomes of farm animals, insects and nematodes (13). At the same time, environmental enterococci with wide-ranging stress resistance offer potential benefits such as tolerance of toxic metals (14) and probiotics (15).

Yet surprisingly little is known about the genetics of enterococcal adaptation to acid or base. In *E. faecium*, acid shock leads to metabolomic changes such as elevated production of dipeptides (16). Acid stress in *Enterococcus* has been studied as part of general stress responses (17–19). At high pH, such as in the presence of the dental application of calcium hydroxide, *Enterococcus* survival requires maintenance of proton motive force (20). Cellular effects of high pH include changes in morphology (21) and in metabolism and protein synthesis (22, 23).

In Gram-negative bacteria, acid adaptation is associated with fitness tradeoffs such as the loss of antibiotic resistance. *Escherichia coli* K-12 experimental evolution at low pH (24, 25) or in the presence of benzoic acid (26, 27) leads to loss of genes encoding multidrug transporters driven by the proton-motive force. It was therefore of interest to explore pH stress adaptation and fitness tradeoffs in a Gram-positive model organism.

We investigated the genetics of pH stress in the model organism *Enterococcus faecalis* OG1RF (2) by means of experimental evolution of planktonic and biofilm cultures. For comparison, in *E. coli*, experimental evolution in acid or base yields mutations in a number of stress response genes, such as amino-acid decarboxylases (24, 25, 28) and acid-resistance regulators (29). In *E. faecalis* pH stress is also important for biofilm formation, including endodontic infections (20, 30). A method to address stress effects in biofilms is experimental evolution by serial bead transfer. Bead-transfer experiments yield informative genetic variants in other environmental bacteria such as *Pseudomonas* and *Burkholderia* species (31, 32). In *Enterococcus* species, experimental evolution has been applied to studies of virulence and antimicrobial susceptibility (33, 34) but not to more fundamental stress factors such as pH.

We find that experimental evolution yields pH-adaptative mutations in a number of genes, notably those encoding elongation factor G (*fusA*) at low pH and the phosphate ABC transporter (*pstABC*) at high pH. Mutations in *fusA* may confer resistance to fusidic acid (35, 36); but our *fusA* mutants show a tradeoff of fusidate sensitivity. The *pts* loci are associated with virulence (37, 38). In our biofilm-evolved strains, base-selected mutations in *pstABC* and in *pyrR* (pyrimidine biosynthesis regulator) led to small-colony growth, a class of phenotype associated with virulence in clinical strains (39, 40).

## RESULTS

### Experimental evolution of acid- and base-adapted mutants

Experimental evolution of *E. faecalis* OG1RF was conducted by microplate serial dilutions at 1:250, as described under Methods. Eight evolving populations were maintained in brain-heart infusion (BHI) buffered at each pH: pH 4.8, pH 7.0, and pH 9.0. After 500 generations, one isolate from each population was resequenced for comparison with the OG1RF reference (2) using the *breseq* resequencing pipeline (41, 42).

The resequenced clones had a range of 1-5 mutations per clone. The mutations predicted were nearly all point mutations or small deletions (Table 1). Only one deletion encompassed more than one gene (genes encoding catechol 2,3 dioxygenase and a multidrug ABC transporter, in clone C06). Overall, the ratio of nonsynonymous to synonymous codon changes was extremely high, with only one silent mutation out of 48. The high proportion of nonsynonymous changes indicates the occurance of adaptive selection (43). Another sign of adaptive selection is that two-thirds of the mutant genes were shared by at least two clones, although the mutations were at different positions. The different pH classes showed a significant difference in mutation frequency:

**Table 1.**
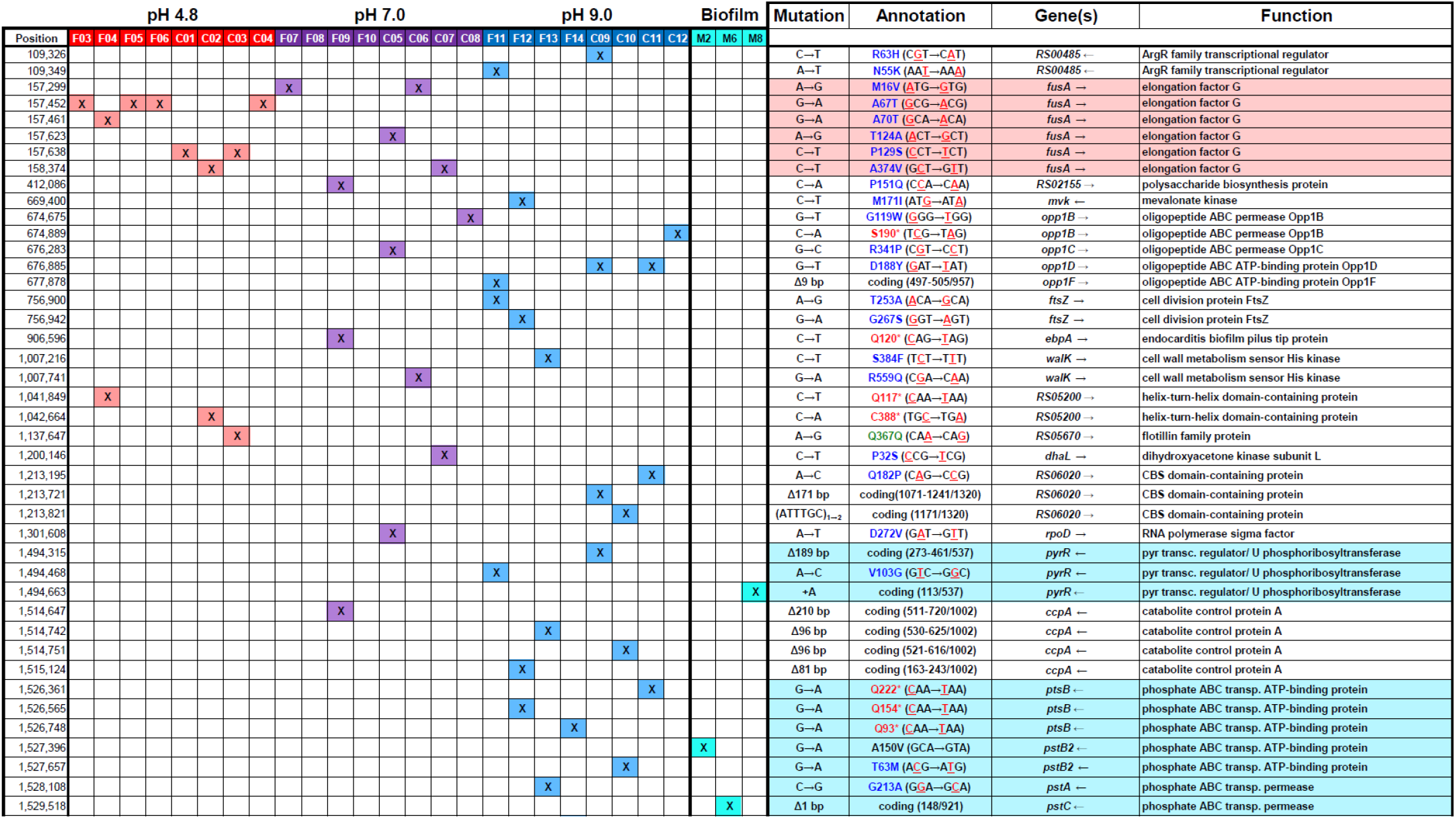
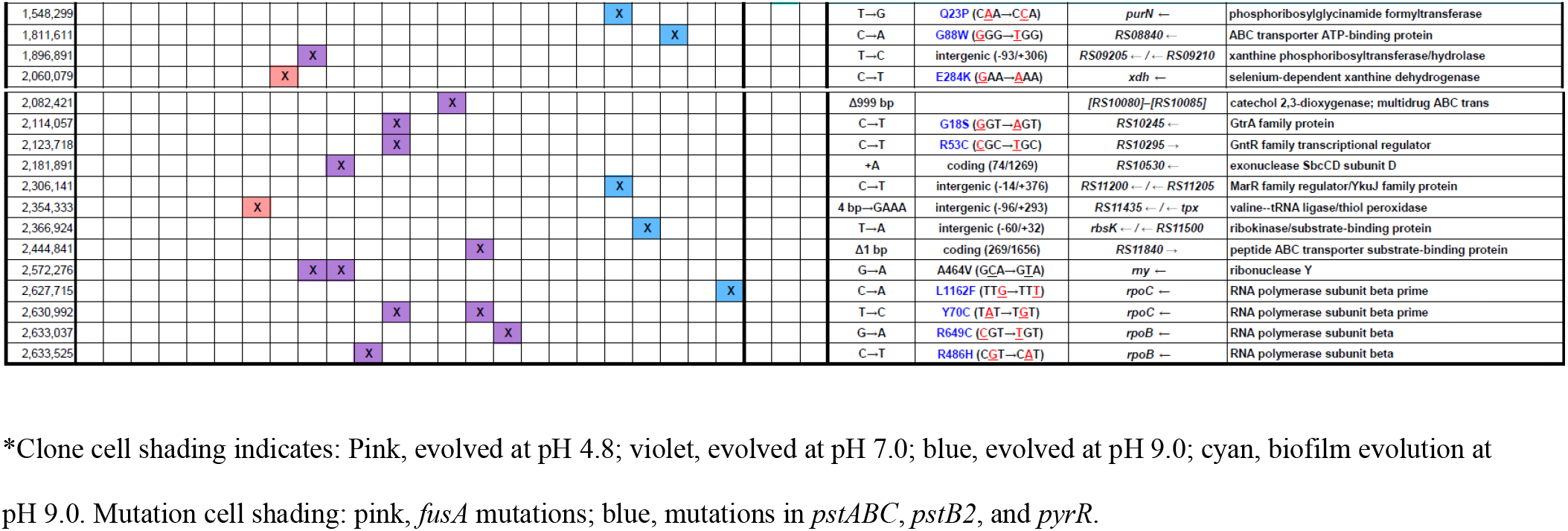
Mutations in clones from experimental evolution*.

### Acid-evolved *fusA* mutants are fusidate-sensitive at pH 7.0

In our acid-evolved clones, the gene that showed the most mutations encodes elongation factor G (*fusA*), an essential translation protein associated with virulence and biofilm formation (44–46). The ancestral strain OG1RF is resistant to fusidic acid, associated with the *fusA* mutation C316A (2). We found an added missense mutation in *fusA* in every acid-evolved clone and in five of our eight pH 7-evolved clones. Six different missense mutations occurred, in different strains. No *fusA* mutations were found in populations evolved at high pH. Two acid-evolved clones contained an additional mutation affecting a gene of unknown function, RS05200, encoding a helix-turn-helix protein. Another clone showed a missense mutation in *xdh* (xanthine dehydrogenase).

The mutant strains from pH 4.8-evolved populations were tested for growth advantage during culture at low pH. (**Fig. 1**). At all three pH levels (pH 4.8, pH 7.0, pH 9.2) the growth curves for mutants were indistinguishable from those of the ancestral strain. For all strains, the steepest rise for log-phase growth occurred at the optimal condition of pH 7. No evidence for directional selection at low pH was seen under the conditions tested.

**Figure 1.**
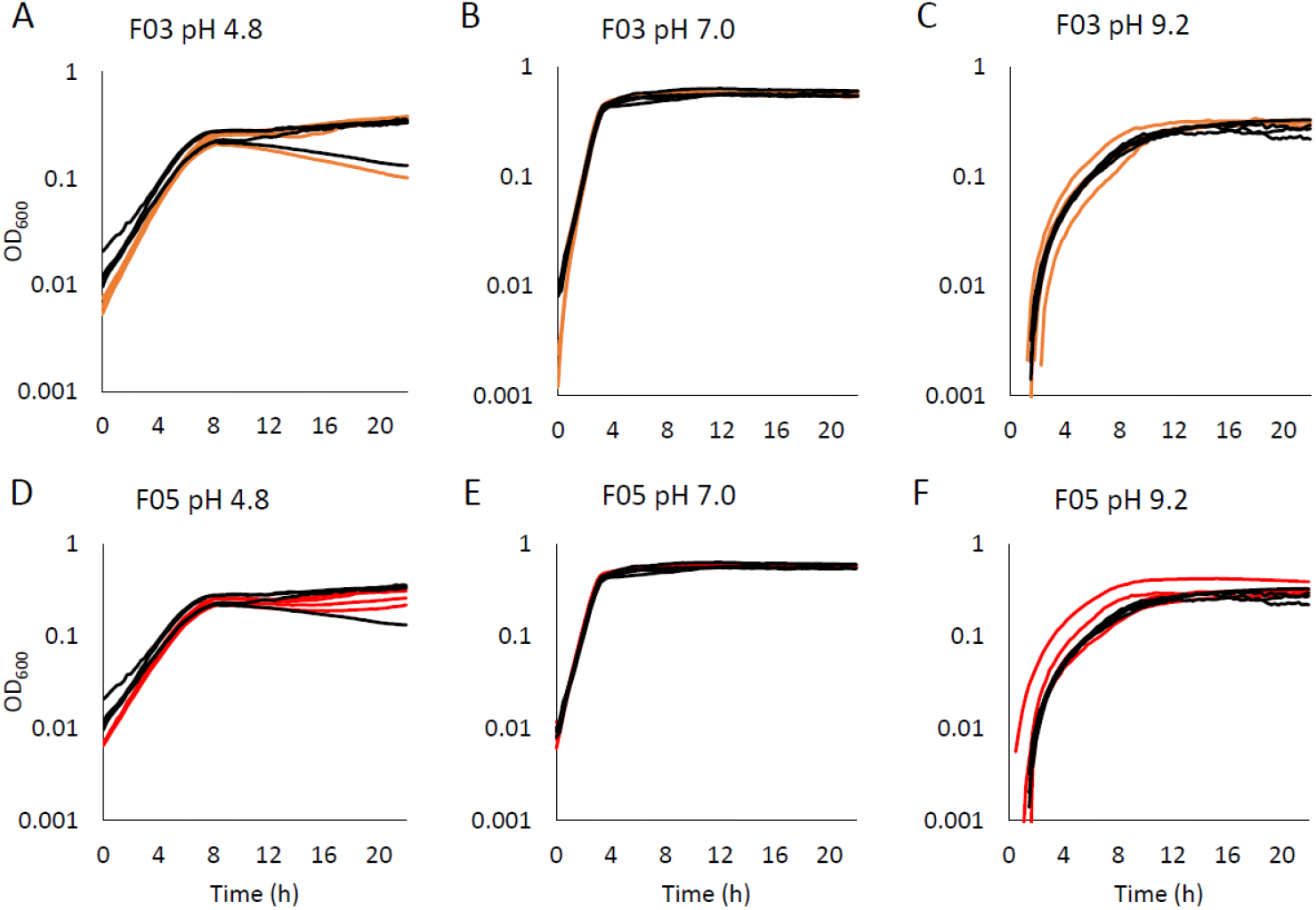
Acid-selected *fusA* mutants show no growth enhancement. *E. faecalis* strains were isolated after 500 generations of planktonic evolution (described under Methods) at pH 4.8. Each clone containing a single mutation in *fusA* was cultured approximately 20 h at 37°C in BHI buffered at pH 4.8, at pH 7.0, or at pH 9.2, then 2 μl was inoculated into 200 μl at the same pH. Four representative curves are shown for each strain: F03 (*fusA*), brown lines (A, B, C); F05 (*fusA*), red lines (D, E, F); OG1RF (ancestor), black lines for all panels.

Since all our acid-evolved clones and four of the eight clones evolved at pH 7 showed additional point mutations in *fusA*, we tested all clones for fusidate resistance in comparison with the ancestor OG1RF (**Fig. 2 and S1**). All clones containing a *fusA* mutation had lower fusidate resistance than OG1RF. However, the clones lacking a *fusA* mutation (F08, F09, F10, C08) showed fusidate resistance indistinguishable from that of the ancestor. Each of the six *fusA* mutations incurred a single amino-acid substitution at sites different from the OG1RF mutation (2). Thus it appears that various kinds of changes in FusA structure can reverse the OG1RF resistance phenotype.

**Figure 2.**
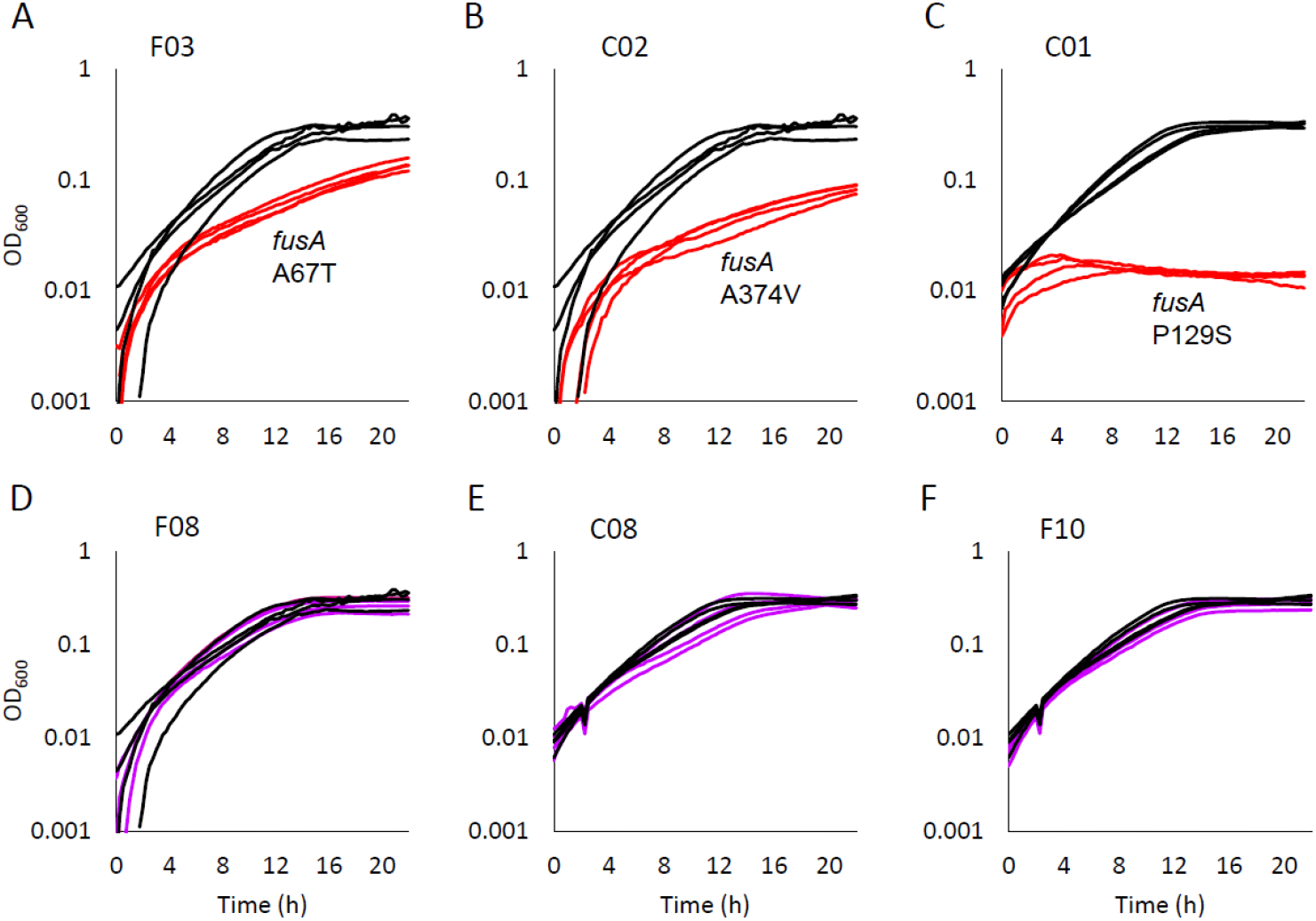
Evolved clones with *fusA* mutations lose resistance to fusidic acid. For each clone, eight replicate cultures were compared with replicates of the ancestor OG1RF. All were cultured with 400 μg/ml fusidic acid, in BHI buffered at pH 7.0 at 37°C. **A, B, C.** Clones evolved at pH 4.8 (red lines) show lower growth than strain OG1RF (black lines). Endpoint growth was compared at 22 h using t-test (p < 0.01). **D, E, F.** Clones evolved at pH 7.0 (purple lines) show growth rates indistinguishable from that of strain OG1RF (black lines).

### Base-evolved populations had mutations in phosphate transport (*pst*) and metabolism (*pyr*)

The base-evolved clones had the largest number of mutations overall (Table 1). All but one of the eight strains from pH 9.0 evolution showed a mutation in a gene involving phosphate metabolism: *pstB, pstB2* (both encoding a phosphate ATP-binding protein of an ABC transporter (47, 48)and *pyrR* (encoding a bifunctional pryimidine transcriptional regulator/ uracil phosphribosyltransferase (49–51)). A majority of these mutations were frameshift or nonsense codons that would knock out the gene. Each strain showed just one mutation in one of these three genes. Several base-evolved clones had additional mutations. Four strains had deletions affecting *ccpA* (catabolite control protein A), and four strains had deletions in various subunits of *oppBCDF* (an oligopeptide ABC transporter). Two strains had missense mutations in an ArgR-family transporter, and another two strains had mutations in septation protein FtsZ.

Several genes were mutated in populations evolved at pH 7.0, as well as in those at pH 9.0. These included *opp1BCDR, ccpA, walK* (cell wall metabolism sensor histidine kinase), and *rpoB, rpoC* (RNA polymerase subunits beta and beta prime). The only gene mutated solely in two pH 7-evolved clones was *rny* (ribonuclease Y).

### Biofilm evolution by bead transfer yields small-colony clones

The evolution experiment was modified to favor genetic variants that adhere to a bead forming a biofilm (31, 32). Cultures underwent four bead transfers over six days, generating approximately 25-50 doublings. The final populations were plated on BHI and inspected for variant phenotypes. No phenotypes were observed after evolution in acid (pH 4.8) or at pH 7, but the pH 9-evolved populations showed two colony size morphs, small and large. Three small-colony clones were found that retained the phenotype upon reculturing: (strains M02, M06, and M08) (**Figure 3)**. The relative size of their colonies compared to those of the ancestor was approximately: M02, 60%; M06, 30% M08, 60%. The genomes of these clones were sequenced, and each revealed a single mutation (Table 1). The mutations affected genes *pstB2* (M02), *pstC* (M06), and *pyrR* (M08).

**Figure 3.**
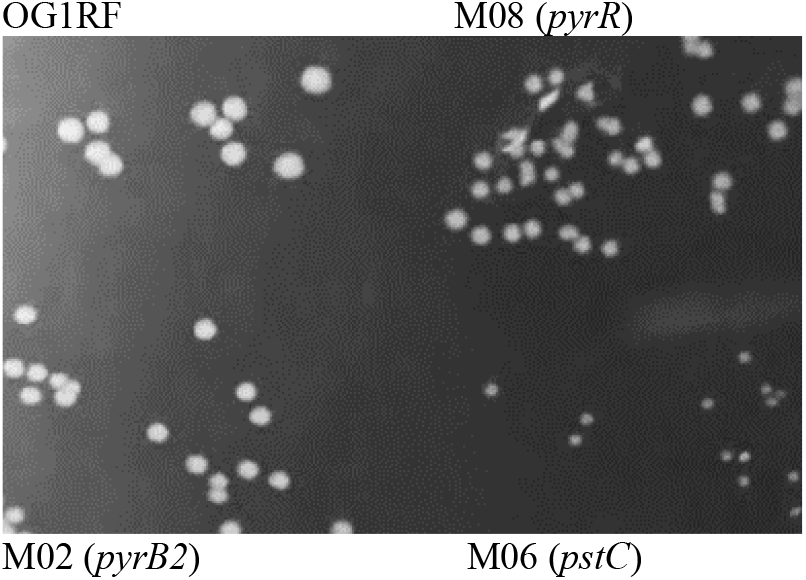
Small-colony phenotype of biofilm-evolved isolates. Each biofilm-evolved strain of *E. faecalis* OG1RF was streaked on BHI and cultured approximately 20 h at 25°C, then approximately 4 h at 37°C.

### Base-evolved clones with *pst* and *pyr* mutations showed evidence of directional selection

We sought evidence for directional selection by observing the batch growth profiles for clones from populations evolved at pH 4.8, pH 7.0 and pH 9.0. Each of the clones was cultured overnight in the three different pH-buffered media, for physiological adaptation to each condition. The strongest evidence for directional selection was observed for pH 9-evolved strains containing a mutation in one of the phosphate-related genes. **Figure 4** shows the growth curves for the biofilm pH 9-evolved strains that each contain one mutation detected by *breseq* in *pstB2* (M02), *pstC* (M06), and *pyrR* (M08). In each case, the log-phase growth (during approximately 0-7 h) was significantly lower at pH 4.8 for the mutant replicates than for replicate curves of the ancestral strain OG1RF; and significantly higher than the ancestor at pH 9.2. (Significance was tested at 6 h, p < 0.05). No consistent selection effect was observed at pH 7.0. These observations are consistent with selection for a high pH-adaptive mutation following prolonged serial culture at high pH, with the directional tradeoff of loss of growth rate at low pH.

**Figure 4.**
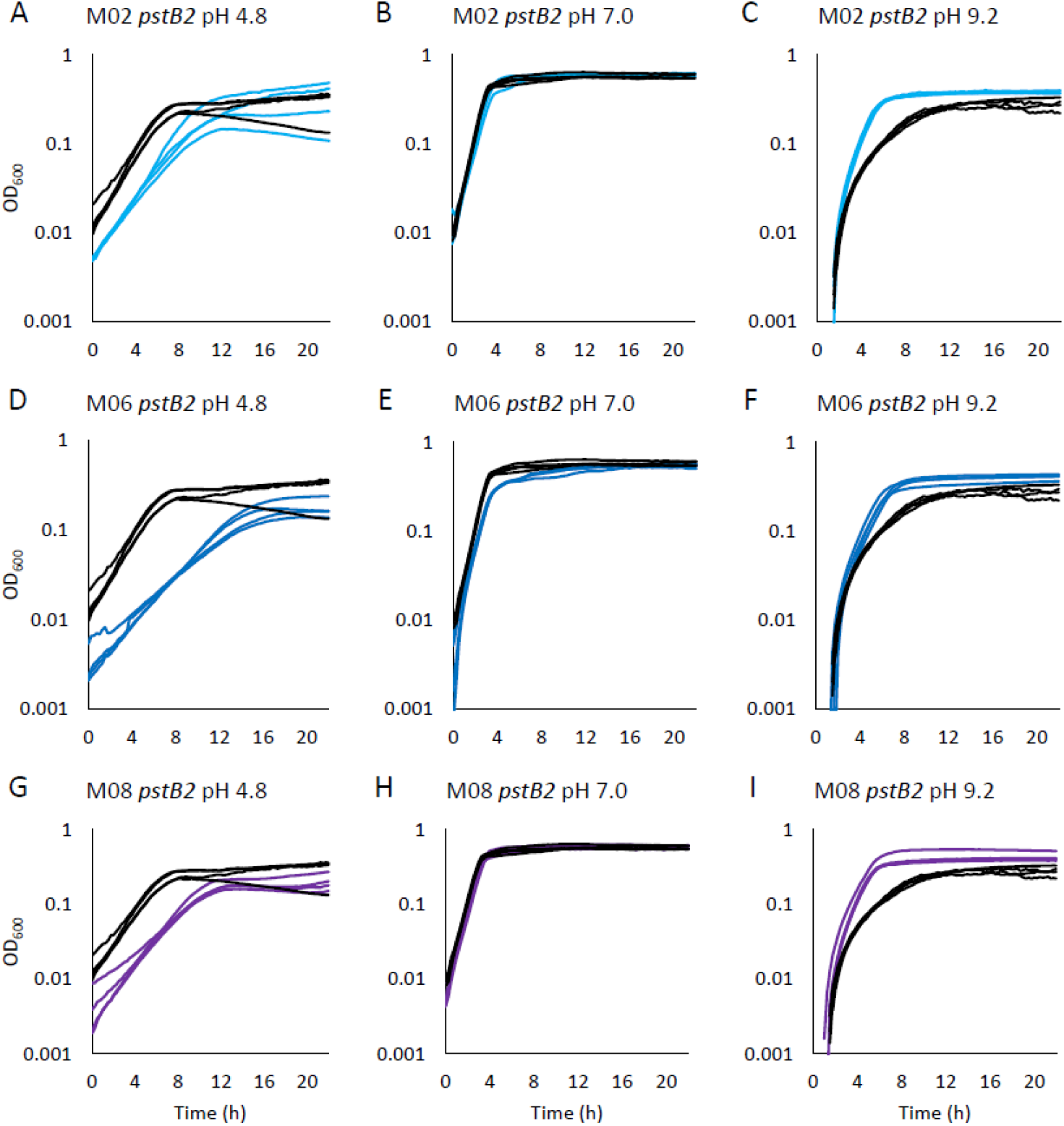
High pH-selected biofilm mutants show increased growth at high pH and decreased growth at low pH. *E. faecalis* strains M02, M06, M08 were isolated as small colonies from bead-transfer biofilm evolution of *E. faecalis* OG1RF at pH 9.0 (described under Methods). Each clone was cultured approximately 20 h at 37°C in BHI buffered at pH 4.8, at pH 7.0, or at pH 9.2, then 2 μl was inoculated into 200 μl of BHI buffered at the same pH. Four representative curves are shown for each strain: M02 (*pstB2*), cyan lines (A, B, C); M06 (*pstC*), blue lines (D, E, F); M08 (*pyrR*), violet lines (G, H, I); OG1RF (ancestor), black lines for all panels. Culture growth was compared at time 6 h by t-test (p < 0.05).

Similar evidence of base-adaptive selection was demonstrated for seven of the eight mutant clones obtained from high-pH planktonic evolution (**Fig. S2**). Only clone C10 showed substantial decrease in growth. Of the eight base-evolved clones, all but clone C12 showed a mutation in *pst* or *pyrR*, as well as mutations in other genes. All eight clones showed a small-colony phenotype on BHI media.

The mutations in phosphate transport favored at high pH might be expected to lead to a dependence on phosphate concentration. However, amendment of the culture media with 10 mM or 50 mM phosphate did not significantly affect the growth curves of evolved strains or of the ancestral strain, either at pH 4.8 or at pH 9.2.

## DISCUSSION

The adaptability of *Enterococcus* bacteria to environmental stresses such as acid or base is important for the human microbiome as well as for soil and plant-associated communities (9, 17–22, 30, 52). Few studies however employ the approach of experimental evolution to examine stress factors under long-term adaptation. Most serial-culture studies of *Enterococcus* sp. focus on virulence or antibiotic resistance rather than environmental factors (34, 53).

We show that experimental evolution of *E. faecalis* leads to distinct pH-dependent patterns of mutations (Table 1). In most cases, genes or operons with variants in multiple clones (that is, the most likely to result from adaptation) are affected only at one end of the pH range; that is, in acid or base but not both. This finding is important as most studies of *Enterococcus* adaptation do not account for pH.

Each of the eight acid-adapted clones had one non-synonymous codon replacement affecting elongation factor G. Three clones from different populations had alanine replaced by threonine; other clones had one of three different substitutions. Four out of eight populations evolving at pH 7 also had a nonsynonymous substitution in *fusA*, but no *fusA* variants appeared at high pH. A possible explanation could be that decrease of cell pH destabilizes the elongation factor G protein, and that the mutations restore protein function. While directional selection for acid adaptation was not shown under the conditions tested, the high frequency of *fusA* missense mutations argues for the occurrence of seletion. Similarly, acid-dependent evolution in *E. coli* leads to mutations in amino-acid decarboxylases known to help pH regulation, although phenotypes are not detected (25).

The *fusA* mutations did show an important tradeoff, the loss of resistance to fusidic acid. This finding implies that *E. faecalis* growth at low pH, such as in dental biofilms and in the intestine (17, 52), could select against some forms of antibiotic resistance. Previously, acid (pH 5.5) increases the fusidate sensitivity of clinical isolates of methicillin-resistant *Staphylococcus aureus* (54). Thus it might be of interest to investigate the role of pH in fusidate therapy.

High pH effects on *Enterococcus* species are important for dental microbiomes because these bacteria can resist endodontic therapies involving application of high-pH pastes such as calcium hydroxide (20, 30). The basis of high-pH resistance is poorly understood. A number of genes are upregulated at high pH (21, 22). One of these, encoding the key septation protein FtsZ, had an amino-acid substitution in two of our high-pH clones (Table 1).

We show evidence for directional selection of variants that knock out a component of the PstABC phosphate uptake system, in most cases by frameshifting deletion or by nonsense substitution (Table 1). Each *pst* mutation we found was associated with higher growth at high pH but lower growth at low pH (Figure 2). The *pstC* mutant (strain M06) showed the smallest colonies; this would make sense if *pstC* knocks out the entire phosphate uptake whereas mutations in the *pstB* or *pstB2* redundent genes only partly cut phosphate uptake.

The growth rate change of *pst* and *pyrR* mutations led to visibly smaller colonies on BHI media. While no change in antibiotic resistance was found under our conditions, in clinical studies small-colony variants are associated with drug resistance and increased virulence (39, 40, 55). Mutations in *pst* genes are associated with clinical virulence (Lieberman) and altered antibiotic resistance (48). In soil and plant root communities, phosphate release is maximized at pH below 7 (56, 57) and has been proposed to contribute to bacterial pH regulation (58). In this model, phosphate release is inhibited at low pH and activated at high pH. Blocking phosphate uptake at high pH could enhance alkali-induced phosphate release.

The effect of *pyrR* loss and it relationship to phosphate is unclear. One possible connection to phosphate transport is the uracil phosporibosyltransferase activity of this regulator (50). This enzyme reaction releases pyrophosphate and has a high pH optimum. Thus it could be that deletion of *pyrR* decreases phosphate release at high pH. In pathogenic staphylococci, *pyrR* is associated with biofilm formation and virulence (51, 59).

Additional mutated genes in high-pH evolved population include two metabolic loci, the oligotransport operon *oppBCD*, and the catabolite reculator *cppA*. Changes in these genes could enhance metabolism under the lowered energy condition of high pH, where the transmembrane pH difference is inverted and the proton potential consists solely of charge difference (60). Overall, our study shows how *E. faecalis* exposure to acid or base can lead to evolutionary shifts with potential importance for host adaptation.

## MATERIALS AND METHODS

### Bacterial strains and culture media

All strains of *E. faecalis* are derivatives of *E. faecalis* OG1RF (2) kindly provided by Barbara Murray. For evolution serial culture, strains were cultured in pH-buffered brain-heart infusion medium under BSL-2 conditions at 25°C. For growth curves, incubation was at 37°C. Buffers for experimental evolution included 100 mM homopiperazine-1,4-bis(2-ethanesulfonic acid) (HOMOPIPES, pK_a_=4.55), and 100 mM N-cyclohexyl-3-aminopropanesulfonic acid (CAPS, pK_a_=10.4). For some of the phenotype testing at pH 7, the buffer included in growth medium was piperazine-N,N’-bis(2-ethanesulfonic acid) (PIPES, pKa=6.8). pH was adjusted with KOH. Fusidic acid was from Millipore-Sigma. Some colony isolations were performed on tryptic soy agar (TSA).

### Experimental evolution of planktonic cells

Serial culture was performed by dilution into deepwell microplates daily, or after two days (every sixth dilution). Eight serial cultures were maintained at each pH: pH 4.8, pH 7.0, and pH 9.0. For each dilution, 2 μL of the previous culture was transferred into 1000 μL (1:250) of fresh medium. All media consisted of brain heart infusion media buffered with 0.1 M each of CAPS HOMOPIPES adjusted with KOH to the appropriate pH. Two days growth was assumed to yield approximately 8 generations (doublings). All evolving populations were propagated to at least 500 generations. From every sixth dilution, 200 μl was combined with 100 μl of 50% glycerol and transferred to a deepwell plate for storage at −80°C.

Once evolved populations reached ~500 generations, each well was streaked on individual plates of brain-heart infusion agar (BHI) and cultured for 48 hours. Two clones were chosen from each well population and used to create individual overnight cultures in BCH. 2 μL of the first set of overnight solutions were used to inoculate 1 ml of fresh BCH overnights. A portion of each clone was converted to freezer stock, and a portion was extracted for genomic DNA.

### Experimental evolution of biofilms

Biofilm evolution was performed by serial bead transfer, based on methods devised by Cooper and colleagues (31, 32). Four replicate populations of *E. faecalis* OG1RF were grown at 25°C in the presence of 7-mm polystyrene beads suspended in 2 ml of BHI buffered with CAPS and HOMOPIPES, adjusted to the appropriate pH. The low pH culture was grown at pH 4.8, while the neutral pH culture was grown at 7.0, and the high pH culture was grown at pH 9.0. Populations were selected for reversible surface attachment through transfer of a bead to a new test tube, in which cells must adhere to a new bead in order to persist, after 48 hours. This process was repeated by alternating white and black beads for four serial dilutions.

At the end of the dilution series, each bead was placed in 1 ml BHI medium to wash off most planktonic cells; then placed again in 1 ml fresh BHI and vortexed for 1 min. The vortexed suspension was then plated on BHI agar plates and incubated 48 h. Plates were then observed for novel colony morphologies.

### Growth Curves

From streaked TSA or BHI plates of ~500-generation evolution isolates, single colonies were picked for overnight culture in BCH buffered at the same pH as the subsequent microplate (pH 4.8, pH 7.0, or pH 9.2). 4 μL of the overnight culture of *E. faecalis* was transferred into sterile 96-well microplates with 200 μL (1:100) of BCH or other media with 100 mM HOMOPIPES and 100 mM CAPS adjusted to the appropriate pH. Growth curves were performed on a Spectramax microplate reader (Molecular Devices) at OD_600_ with OD reads taken every 15 minutes for 22 hours. Comparison of batch growth was performed by T-test at standard intervals post inoculation.

### DNA sequence analysis

For the planktonic evolution cultures, DNA was extracted from one selected ~500-generation isolate of each well (24 isolates) and from the OG1RF(B) ancestral strain (2 isolates). The 24 sequenced clones were designated BF01-14 and C01-12. For the biofilm evolution at pH 9, the final biofilm bead suspensions were streaked on BHI agar. After 48 hours, one large colony and one small colony variant (SCV) were isolated from each plate and streaked on new BHI plates separately. The isolates were plated on BHI twice more to ensure the persistence of the growth phenotype. Three large-colony isolates (M1, M5, M7) and three small-colony isolates (M2, M6, M8), were selected for DNA preparation and sequencing.

Genomic DNA was extracted using the MasterPure Gram Positive DNA Purification Kit. Purity and concentrations of DNA were determined using a NanoDrop 2000 spectrophotometer (Thermo Fisher Scientific). DNA was sequenced by Admera Health (South Plainfield, NJ). Library preparation was performed using NexteraXT library kit (Illumina, California, USA). Sequencing was performed via Illumina NovaSeq (Illumina, California, USA) with a read length of 150 PE for 40 M PE reads.

Mutations were called from the DNA sequences by alignment to the *E. faecalis* OG1RF reference sequence with NCBI accession number NC_017316.1 using the *breseq* version 0.36.1 resquencing pipeline (41, 42). Resulting genomic data had approximately 300-fold coverage. Our lab stock of OG1RF (the ancestor for all evolution experiments) showed two differences from the reference that appeared in all our evolved clones: insertion of A at genome position 260,670, causing frameshift in gene RS01405 (acyl-ACP thioesterase); and T deletion at position 2,108,103 within an intergenic region.

## Data availability

All sequence data are deposited at NCBI, SRA PRJNA933188. Reviewer link: https://dataview.ncbi.nlm.nih.gov/object/PRJNA933188?reviewer=h51sokrkgscppqt0618086ihqj

## AUTHOR CONTRIBUTIONS

BAF performed the planktonic evolution, and AW performed the biofilm evolution; they both drafted a manuscript. BAF, AW and ZS conducted growth curves. JLS conceived the study, mentored the students and completed the manuscript.

## ACKNOWLEDGMENTS

This work was supported by National Science Foundation award MCB-1923077. B. F. and A. W. designed the study, conducted experiments and drafted a manuscript. Z. S. conducted experiments and analysis. J. S. conceived the study, mentored the students and completed the manuscript for publication. We thank Daniel Barich for assistance with computing and statistics.

