## Supplemental Figures for "*Enterococcus faecalis* OG1RF Evolution at Low pH Selects Fusidate-sensitive Mutants in Elongation Factor G and at High pH Selects Defects in Phosphate Transport"

For

By

Bailey A. Fitzgerald,[1] Ayman Wadud,[1] Zachary Slimak, and Joan L. Slonczewski*

*Corresponding author: Joan L. Slonczewski, Kenyon College, Gambier, OH 43022

[1]These two authors contributed equally to this study.

Figure S1. Fusidate sensitivity of clones evolved at pH 4.8 (A-H) or at pH 7.0 (I-P).

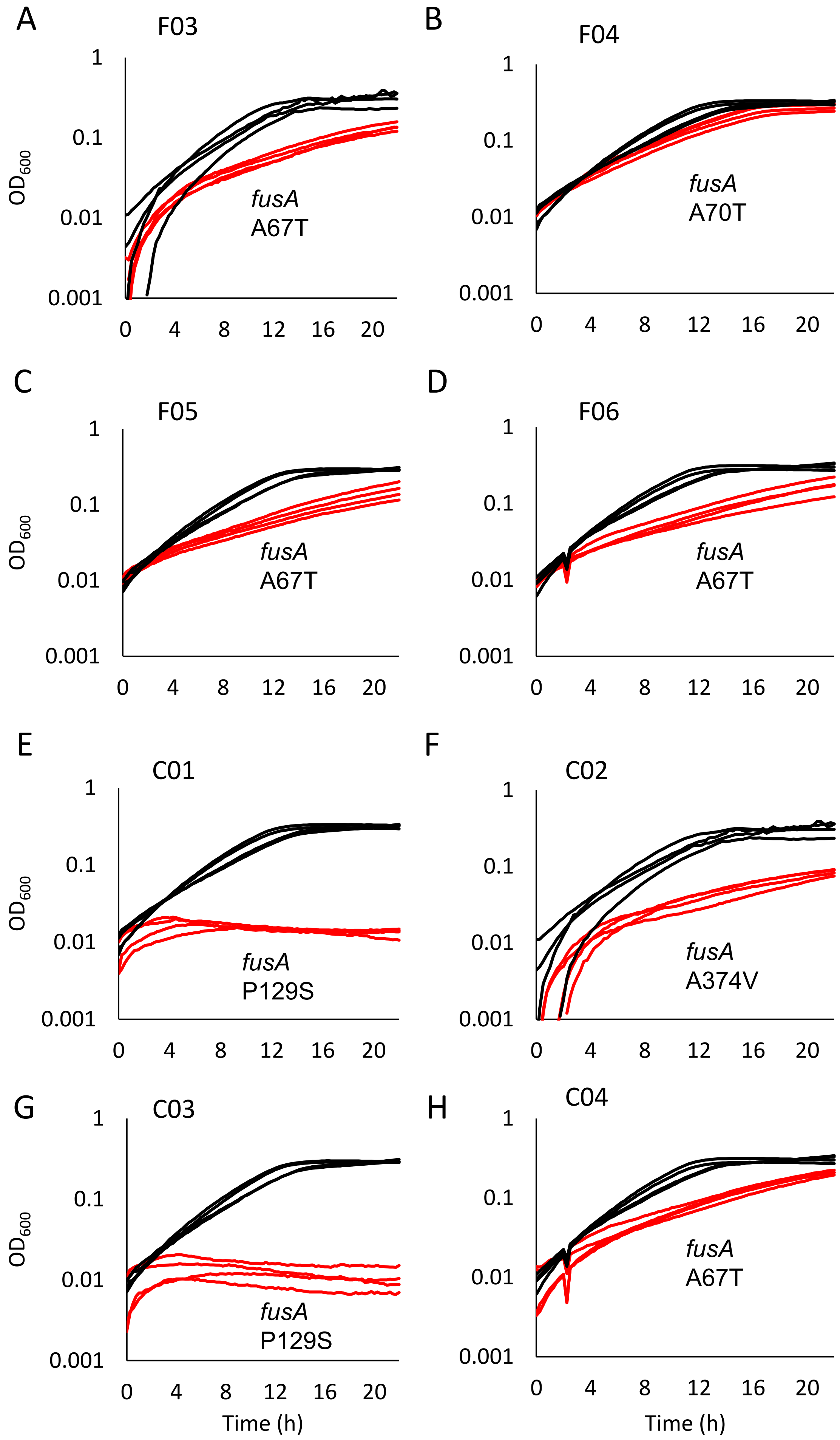

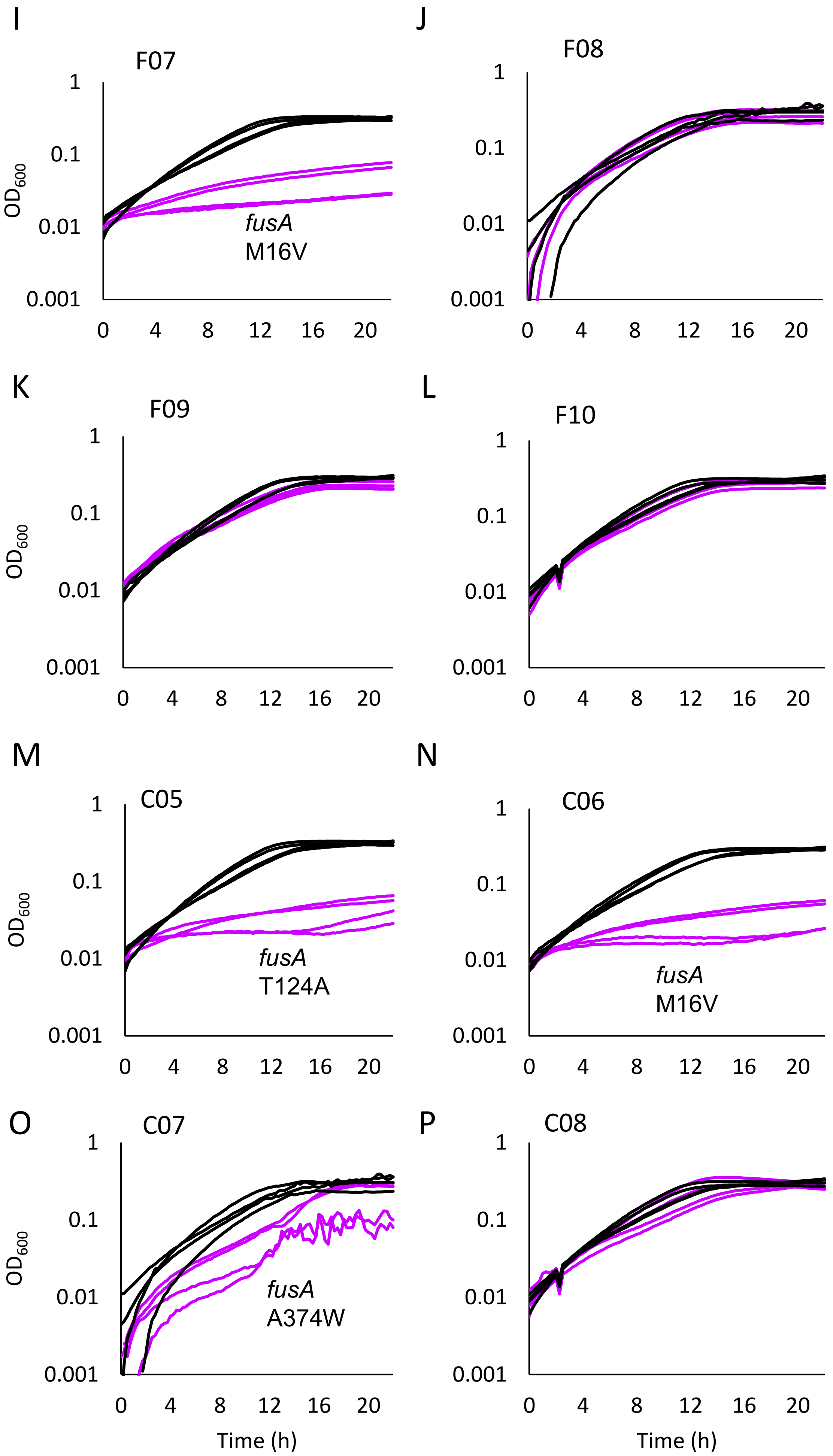

I
F07
OD600
fusA
M16V
J
F08
K
F09
L
F10
M
C05
fusA
T124A
N
C06
fusA
M16V
O
C07
fusA
A374W
P
C08
Time (h)
Time (h)
1
0.1
0.01
0.001
0
4
8
12
16
20

**Figure S1. Fusidate sensitivity of clones evolved at pH 4.8 (panels A-H) or at pH 7.0 (panels I-P).** For each clone, eight replicate cultures are compared with replicates of the ancestor OG1RF. All were cultured in BHI with 400 µg/ml fusidic acid, buffered at pH 7, as described for Figure 2. The $OD_{600}$ culture values at 22 h for each clone versus OG1RF were compared using t-test.

t-test p-values:
A = 0.0074
B = 0.049
C = 0.0031
D = 0.0023
E = 5.3E-05
F = 0.0041
G = 4.2E-06
H = 0.0042
I = 5.4E-06
J = 0.30
K = 0.0045
L = 0.24
M = 6.9E-07
N = 2.8E-06
O = 0.106
P = 0.36

Figure S2. High pH-selected clones cultured at pH 4.8 and at pH 9.2.

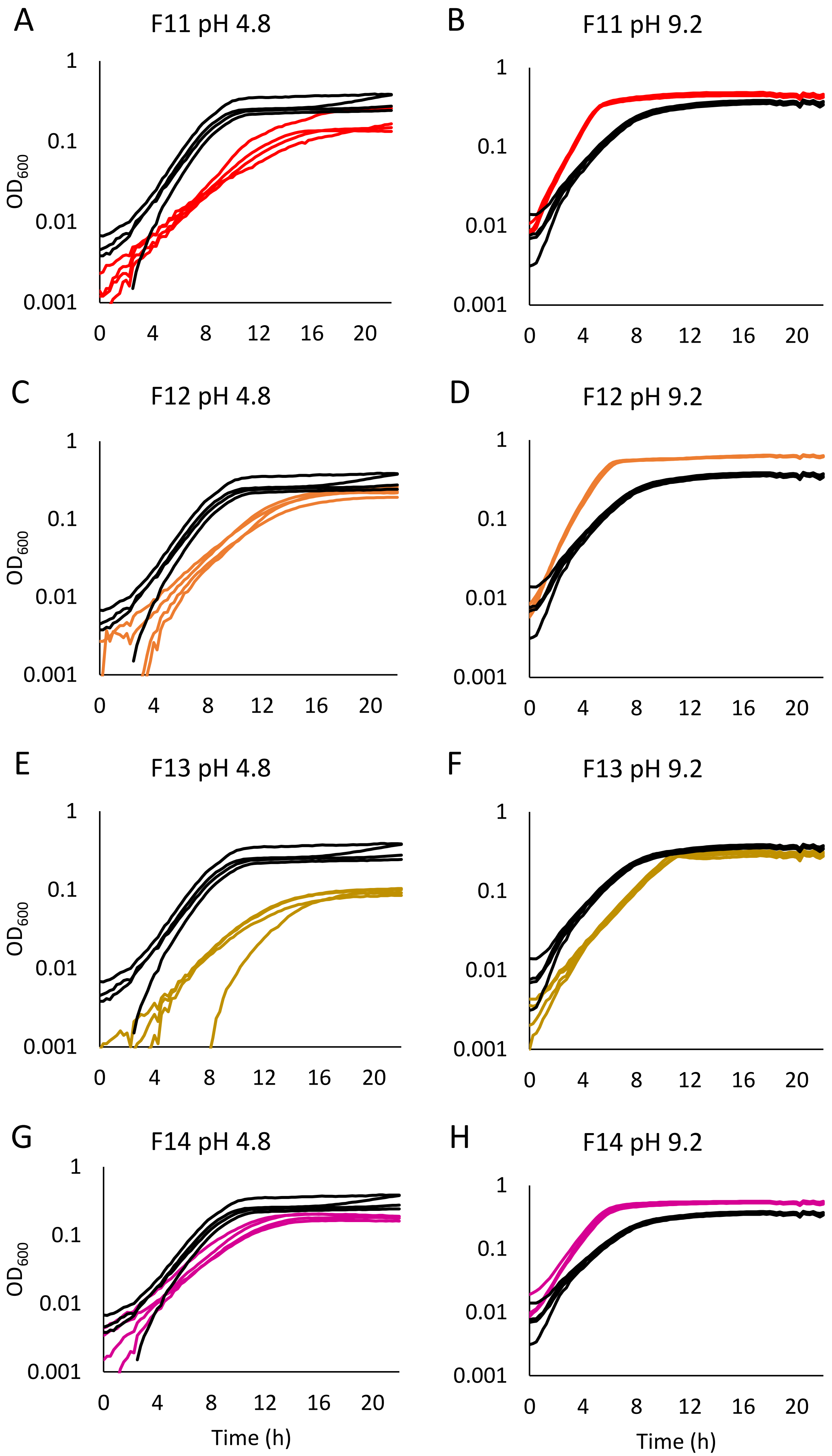

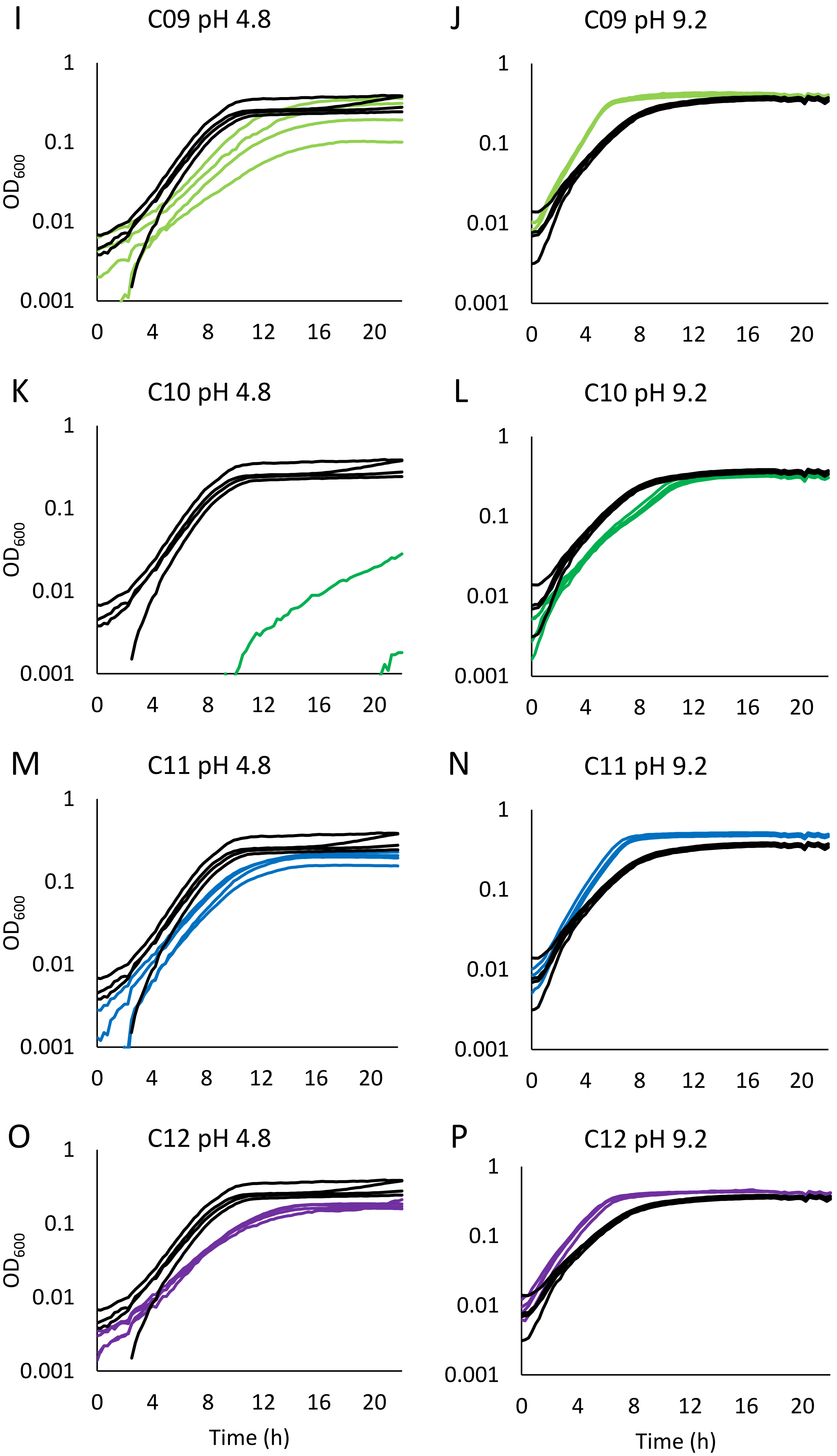

I
C09 pH 4.8
J
C09 pH 9.2
K
C10 pH 4.8
L
C10 pH 9.2
M
C11 pH 4.8
N
C11 pH 9.2
O
C12 pH 4.8
P
C12 pH 9.2
OD600
Time (h)
1
0.1
0.01
0.001
0
4
8
12
16
20

**Figure S2. High pH-selected clones cultured at pH 4.8 and at pH 9.2.** For each clone, eight replicate cultures are compared with replicates of the ancestor OG1RF. Panels A, C, E, G, I, K, M, O – strains were cultured at pH 4.8; panels B, D, F, H, J, L, M, P – strains were cultured at pH 9.2. Culture conditions were as for Figure 4. The $OD_{600}$ culture values at 6 h for each clone versus OG1RF were compared using t-test.

t-test p-values:
A = 0.021
B = 0.00069
C = 0.067
D = 1.67E-06
E = 0.0076
F = 0.0015
G = 0.026
H = 3.0E-05
I = 0.282
J = 0.0116
K = 0.0025
L = 0.0069
M = 0.0327
N = 6.0E-05
O = 0.026
P = 0.0042
